# Effect of high-frequency low-intensity pulsed electric field on protecting SH-SY5Y cells against hydrogen peroxide and β-amyloid-induced cell injury via ERK pathway

**DOI:** 10.1101/2019.12.30.891499

**Authors:** Wei-Ting Chen, Guan-Bo Lin, Yu-Yi Kuo, Chih-Hsiung Hsieh, Chueh-Hsuan Lu, Yi-Kun Sun, Chih-Yu Chao

## Abstract

As the most common type of neurodegenerative diseases (NDDs), Alzheimer’s disease (AD) is thought to be caused mainly by the excessive aggregation of β-amyloid protein (Aβ). However, a growing number of studies have found that reactive oxygen species (ROS) play a key role in the onset and progression of AD. The present study aimed to probe the neuroprotective effect of high-frequency low-intensity pulsed electric field (H-LIPEF) for SH-SY5Y cells against hydrogen peroxide (H_2_O_2_) and Aβ-induced cytotoxicity. By looking in a systematic way into the frequency- and amplitude-dependent neuroprotective effect of pulsed electric field (PEF), the study finds that H-LIPEF at 200 Hz produces the optimal protective effect for SH-SY5Y cells. The underlying mechanisms were confirmed to be due to the activation of extracellular signal-regulated kinase (ERK) pathway and the downstream prosurvival and antioxidant proteins. Because the electric field can be modified to focus on specific area in a non-contact manner, the study suggests that H-LIPEF holds great potential for treating NDDs, whose effect can be further augmented with the administering of drugs or natural compounds at the same time.

## Introduction

Neurodegenerative diseases (NDDs) feature progressive deterioration of brain function, resulting from the gradual degeneration or death of neurons. Along with increasing human lifespan, deaths caused by NDDs have been on the rise over the past years, impacting the quality of life of patients and their families. The World Health Organization (WHO) predicts that by 2040, NDDs will overtake cancer to be the second leading cause of death, next to cardiovascular diseases.

As the most common kind of neurodegenerative disorder among various NDDs, Alzheimer’s disease (AD) displays such typical pathology including the deposition of extracellular cortical plaques formed by aggregation of β-amyloid protein (Aβ) and intracellular neurofibrillary tangles caused by hyperphosphorylation of Tau protein [1]. Scientists have long suspected that these abnormal structures are major culprit in damaging and killing nerve cells, which causes progressive brain degeneration and deterioration of cognitive function among elderly people. Anti-amyloid therapies have long been pursued by scientists, however, a more nuanced treatment is required because of the continuing clinical failures [2]. In addition, there has been increasing evidence showing that ROS plays a central role in the onset and progression of AD [3]. Therefore, drugs or treatments which could decrease the ROS level and protect neural cells against oxidative damage may be potential strategies to treat AD. Some in vitro studies have explored the therapeutic effect of antioxidant and antiapoptotic drugs in AD treatment [4, 5]. Clinical trials have been conducted to search for antioxidant supplements or drugs to treat AD but the results have been disappointing [6-10]. There has yet to be an effective treatment for AD or other kinds of NDDs. Existing medications can only avoid deterioration of symptoms or postpone their appearance at best. One blockade in AD drug development is the blood-brain barrier (BBB), which is insurmountable for 98% of small molecule drugs and 100% of large molecule drugs [11]. Therefore, effective AD treatment remains a major challenge for modern medicine.

In our previous study, non-invasive low-intensity pulsed electric field (LIPEF) was employed, in conjunction with fucoidan, in preventing oxidative stress-induced mouse motor neuron death [12]. It found that LIPEF suppressed the hydrogen peroxide (H_2_O_2_)-enhanced expression of ROCK protein and increased the phosphorylation of Akt in H_2_O_2_-treated NSC-34 cell line. The current study employed non-contact LIPEF stimulation in studying human neural cell line SH-SY5Y, a neurodegenerative damage model with extensive application, and probed the neuroprotective effect of LIPEF against oxidative damage caused by H_2_O_2._ It also applied AD disease model by Aβ protein administration in neuroprotective study. Interestingly, the study finds that the LIPEF parameters applied in mouse motor neuron are not effective in human neural cell line. The systematic study on frequency- and amplitude-dependent neuroprotective effect of LIPEF was performed, and the result found that high-frequency LIPEF (H-LIPEF) at 200 Hz had the optimal protective effect for SH-SY5Y cells. Since the alternating current (AC) signal has only one frequency and intensity, it needs time-consuming and painstaking effort for finding the specific AC electric signal to achieve a certain biological effect. Therefore, the approach has only limited applicability, as the specific AC signal is by no means sufficient in targeting complex signal transductions in a biological process or disease treatment. By contrast, the pulsed electric field (PEF) features a composition of multiple sinusoidal subcomponents with diverse frequencies and intensities simultaneously [12]. The study points out that thanks to its multi-frequency components, PEF can facilitate significantly the development of new disease therapeutics in fulfilling different demands.

In the study, we applied the H-LIPEF stimulation to human neural cell line SH-SY5Y, and examined the prosurvival effect of such physical stimulation against oxidative damage caused by H_2_O_2_ and Aβ-induced cytotoxicity. The results found that exposure of the cells to H-LIPEF at 200 Hz significantly ameliorated the H_2_O_2_- and Aβ-induced cytotoxicity in SH-SY5Y cells. Examination of the underlying mechanism also showed that 200 Hz H-LIPEF could activate extracellular signal-regulated kinase (ERK) pathway and downstream neuroprotective proteins. These findings imply that H-LIPEF is a promising electrotherapy, shedding light on a novel approach in treatment of AD or other NDDs.

## Materials and methods

### Experimental setup for exposure of the cells to non-contact H-LIPEF

The LIPEF device described previously [13] was used for exposure of electric field with various strengths and frequencies. The cells were placed between two copper flat and parallel electrodes. Consecutive pulses with pulse duration 2 ms, different electric field intensities and different frequencies were applied across the electrodes. Cells treated with continuous LIPEF were kept in a humidified incubator of 5% CO_2_ and 95% air at 37°C. The control cells were incubated in a similar LIPEF device without the electric field turned on. After optimization of the pulse parameters, we used the H-LIPEF stimulation to produce the electric field in a non-contact manner, and examined its neuroprotection effect against the oxidative insult caused by H_2_O_2_ to SH-SY5Y human neural cells.

### Cell culture and treatment

The SH-SY5Y cells were purchased from Bioresource Collection and Research Center (Hsinchu, Taiwan) and maintained in MEM/F-12 mixture containing 10% fetal bovine serum (FBS) (HyClone; GE Healthcare Life Sciences) and 1% penicillin-streptomycin, supplemented with 1 mM sodium pyruvate and 0.1 mM non-essential amino acids. Cells were maintained in a humidified incubator composed of 5% CO_2_ and 95% air at 37°C. Cells were harvested with 0.05% trypsin–0.5 mM EDTA solution (Gibco; Thermo Fisher Scientific, Inc.) and prepared for in vitro experiments. To differentiate SH-SY5Y cells, the cells were incubated in low serum medium (0.5% FBS) containing 20 µM retinoic acid (RA) (Sigma-Aldrich; Merck KGaA) for 5 days. Cells were pretreated with H-LIPEF for 4 h and then challenged with 500 µM H_2_O_2_ [14, 15] in the continuous administration of H-LIPEF for another 24 h. For the AD disease model, the cells were pretreated with H-LIPEF for 4 h and then challenged with 25 or 50 µM Aβ_25-35_ protein solution [16, 17] in the continuous administration of H-LIPEF for another 4 days. The Aβ stock solution was prepared by solubilizing the Aβ_25-35_ peptide (Sigma-Aldrich; Merck KGaA) in sterile deionized water to a concentration of 1 mM and then incubated at 37°C for 3 days to allow self-aggregation before treatment. After the treatment, the cell viability was determined by MTT assay. Aβ fibrilization was determined by Thioflavin T (ThT) (Sigma-Aldrich; Merck KGaA) staining. Aβ stock solution was diluted with PBS to 25 µM Aβ solution containing 10 µM ThT. The ThT fluorescence was then measured by SpectraMax i3x plate reader (Molecular Devices, LLC.) with excitation filter of 450 nm and emission filter of 490 nm. For the ERK inhibitor experiment, 50 µM PD98059 (Cell Signaling Technology, Inc.) was treated 1 h before the H-LIPEF treatment.

### Cell viability assay

Viability of SH-SY5Y cells after treatments with or without H-LIPEF was assessed by 3-(4,5-dimethylthiazol-2-yl)-2,5-diphenyltetrazolium bromide (MTT) (Sigma-Aldrich; Merck KGaA) assay. Briefly, the medium was replaced with MTT solution (0.5 mg/mL in SH-SY5Y culture medium) and incubated at 37°C for 4 h. During this period, the ability of mitochondrial dehydrogenases to reduce the MTT to formazan crystal is an indicator of cellular viability. Following MTT reaction, formazan dissolution was performed by using 10% sodium dodecyl sulfate (SDS) and 0.01 M HCl. The optical density of each well was then determined at 570 nm subtracting the background at 690 nm using an ELISA microplate reader. The cell viability was calculated based on the intensity of formazan, and expressed as percentage of the non-treated control.

### Flow cytometric detection of apoptotic cells

The mode of cell death was analyzed by flow cytometry with an Annexin V-FITC and propidium iodide (PI) double-staining kit (BD Biosciences). After the treatments, SH-SY5Y cells were collected by trypsinization and resuspended in binding buffer containing Annexin V and PI. Cells were incubated for 15 min at 25°C in the dark before being analyzed by flow cytometry.

### ROS level detection

After the treatments, the SH-SY5Y cells were collected and the ROS levels were detected using the fluorescent dye dihydroethidium (DHE) (Sigma-Aldrich; Merck KGaA). Cells were incubated with 5 µM DHE dye for 20 min at 37°C in the dark and then the fluorescence intensity was measured by flow cytometry in the PE channel. The ROS levels were expressed as mean fluorescence intensity for comparison.

### Mitochondrial membrane potential (MMP) measurement

The detection of MMP was carried out by flow cytometry using 3,3’-dihexyloxacarbocyanine iodide (DiOC_6_(3)) (Enzo Life Sciences International Inc.). DiOC_6_(3) is a lipophilic cationic fluorescent dye which is selective for the mitochondria of live cells. It allows estimation of the percentage of cells with depleted MMP which reflects the initiation of proapoptotic signal. Cells treated with H_2_O_2_ and H-LIPEF alone or in combination for 24 h were harvested and suspended in PBS with 20 nM DiOC_6_(3). After incubation at 37°C for 15 min in the dark, the fluorescence intensity was immediately measured by flow cytometry.

### Western blot analysis

After the treatments, cells were washed with phosphate buffered saline (PBS) and then harvested and lysed in RIPA lysis buffer (EMD Millipore) containing fresh protease and phosphatase inhibitor cocktail (EMD Millipore). The lysates were then centrifuged, and the supernatant was collected and used for the determination of protein concentration. Equal amounts of protein extracts were loaded in the 10% SDS-PAGE wells and transferred to polyvinylidene fluoride membranes. After drying for 1 h at room temperature (RT), the membranes were probed with ROCK (Abcam, plc.), phosphorylated ERK (p-ERK), total ERK (t-ERK), Nrf2, phosphorylated CREB (p-CREB), total CREB (t-CREB), Bcl-2, Bax (Cell Signaling Technology, Inc.), and GAPDH (GeneTex, Inc.) antibodies overnight at 4°C. Subsequently, the washed membranes were incubated with horseradish peroxidase-coupled secondary antibodies (Jackson ImmunoResearch Laboratories, Inc.) for 1 h at RT. All the antibodies were diluted at the optimal concentration according to the manufacturer’s instructions. Finally, protein bands were visualized with an enhanced chemiluminescence substrate (Advansta, Inc.) and detected with the Amersham Imager 600 imaging system (GE Healthcare Life Sciences). The images were analyzed with Image Lab software (Bio-Rad Laboratories, Inc.). For normalization of proteins, GAPDH was used as an internal control.

### Statistical analysis

Each data point represents the average from larger or equal to three independent experiments and is expressed as the mean ± standard deviation. Statistical analyses were performed using OriginPro 2015 software (OriginLab). Differences of statistical significance were analyzed by one-way or two-way analysis of variance (ANOVA), followed by Tukey’s post-hoc test. P-value < 0.05 was considered to indicate a statistically significant difference.

## Results

### In vitro-applied H-LIPEF

The H-LIPEF was applied in a non-contact manner with the apparatus consisting of two thin parallel conductive electrodes mounted on the insulating acrylic frame (Fig 1A). The culture dishes were placed between the parallel electrodes so the cells were not in contact with the electrodes directly. Therefore, the H-LIPEF stimulation device can provide the electric field in a non-invasive manner, avoiding the harmful effects caused by conduction current and Joule effect via contact electrodes. The pulsed electric signal was from a function generator and amplified by a voltage amplifier. In the study, the applied electric signal was characterized by a pulse train waveform (Fig 1B) with various field intensities (1-60 V/cm) and frequencies (2-499 Hz) for optimization of the treatment.

**Fig 1.**
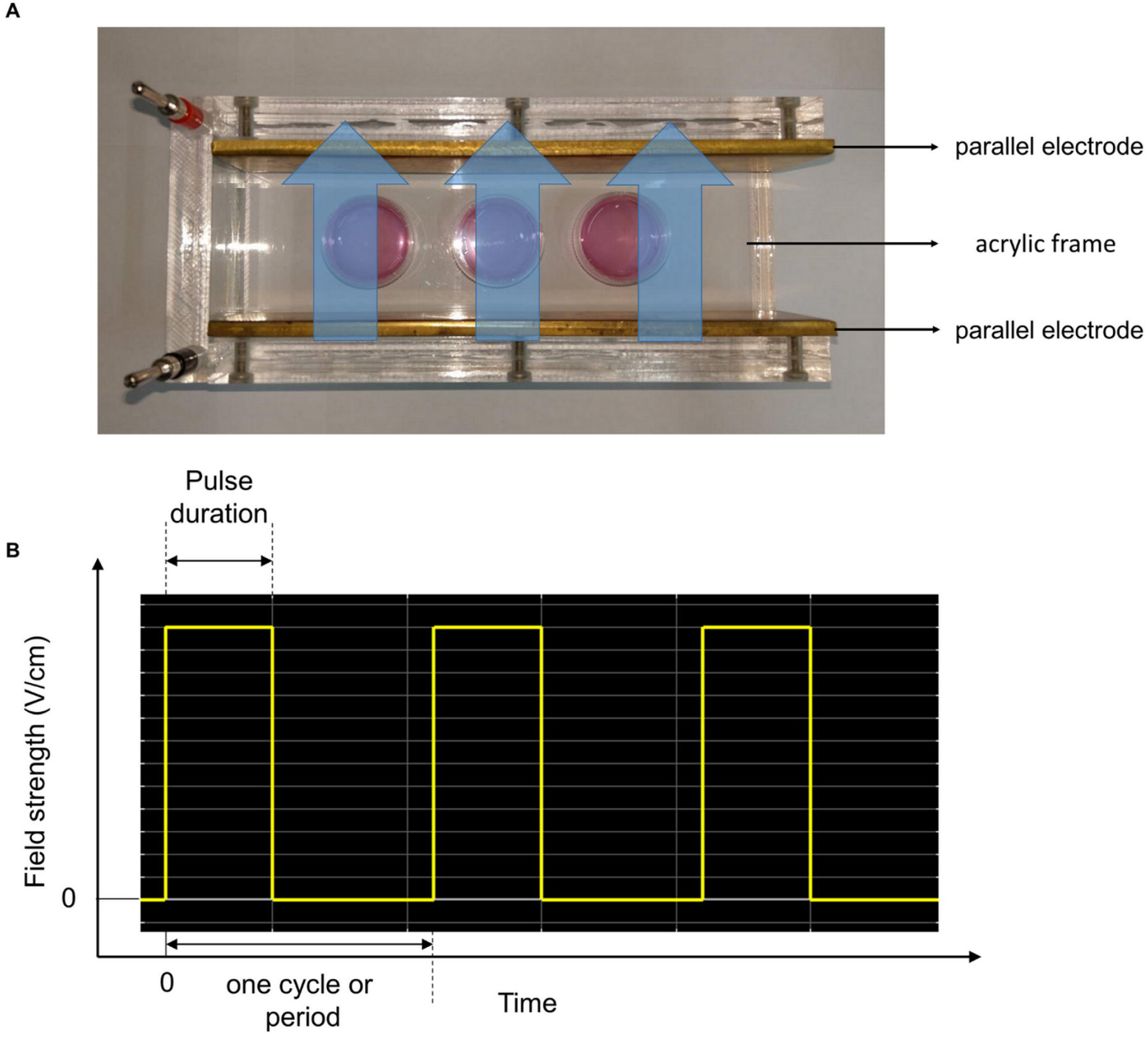
The experimental setup of H-LIPEF. (A) Image of the H-LIPEF stimulation device. The culture dishes were placed in the stimulation device constructed of two parallel electrodes. The blue arrows shown here indicate the schematic representation of non-invasive H-LIPEF exposure on cells. (B) Schematic representation of the pulse train waveform applied on the electrodes.

### Effect of H-LIPEF exposure on H_2_O_2_-induced cytotoxicity in SH-SY5Y cells

The oxidative stress plays a critical role in the etiology and exacerbates the NDDs progression. We applied H_2_O_2_ to SH-SY5Y cells as an oxidative stress and looked into the neuroprotective effect of H-LIPEF technique. As shown in Fig 2A, administration of H_2_O_2_ on SH-SY5Y cells for 24 h decreased the cell viability in a concentration-dependent manner. Treatment of SH-SY5Y cells with 500 µM H_2_O_2_ for 24 h reduced the cell viability to 52% compared to the control cells, which was used in the subsequent experiments. To examine whether H-LIPEF could produce protection and increase the cell viability against the oxidative insult, cells were pretreated with H-LIPEF for 4 h and then exposed to H_2_O_2_ in the continuous administration of H-LIPEF for another 24 h. In our previous study, the low frequency LIPEF with parameters of 2 Hz, 60V/cm, and 2 ms pulse duration was used in the mouse motor neurons NSC-34 [12], but the result showed that this parameter combination is ineffective in the SH-SY5Y cells. In this study, we used H-LIPEF in our experiments, and found that the H-LIPEF stimulation using the parameters of 200 Hz, 10 V/cm, and 2 ms pulse duration produced significant protection against the oxidative insult caused by H_2_O_2_ to SH-SY5Y human neural cell line. To examine whether the frequency and intensity are both critical parameters in the neuroprotective effect, the systematic study on frequency- and amplitude-dependent effects was performed. The intensity of H-LIPEF was firstly fixed at 10 V/cm and the frequency was varied from 2 Hz to 499 Hz. The upper limit was set to 499 Hz since for 2 ms pulse duration, two pulses will overlap for frequency greater or equal to 500 Hz. The results showed that for frequency less than or equal to 100 Hz, their survival rates were not significantly different from the H_2_O_2_-treated group. At high frequency region, the H-LIPEF conferred protective effect to the H_2_O_2_-treated SH-SY5Y cells, and the maximum protective effect was at near 200 Hz (Fig 2B). For the intensity-dependent effect, the frequency was fixed at 200 Hz, and the intensity varied from 1∼60 V/cm was examined. As shown in Fig 2C, it was obvious that there was a maximum at 10 V/cm, indicating that the best protective effect was at 10 V/cm. Therefore, we conclude that both the frequency and intensity are critical parameters in the neuroprotective effect, and the optimum protective parameters (200 Hz, 10 V/cm, and 2 ms pulse duration) (Fig 2D) were used in the following experiments. To further confirm the mode of cell death induced by H_2_O_2_, cells were stained with Annexin V-FITC and PI and assessed by flow cytometry. The results showed that H_2_O_2_-induced apoptotic cells were significantly reduced under H-LIPEF treatment (Fig 3), indicating SH-SY5Y cells were undergoing apoptotic cell death when treating with H_2_O_2_, and H-LIPEF could ameliorate the toxic effect. SH-SY5Y cell is a human neuroblastoma cell line, which is widely used as a neuronal cell model in studying NDDs. To investigate whether H-LIPEF could also provide neuroprotective effect in differentiated SH-SY5Y cells, SH-SY5Y cells were differentiated with RA for 5 days prior to H_2_O_2_ or H-LIPEF treatment. As shown in Fig 4A, the differentiated SH-SY5Y cells exhibited neuronal-like morphology with high levels of neurite outgrowth. And the results showed that H-LIPEF could provide similar neuroprotective effect in the differentiated SH-SY5Y cells (Fig 4B).

**Fig 2.**
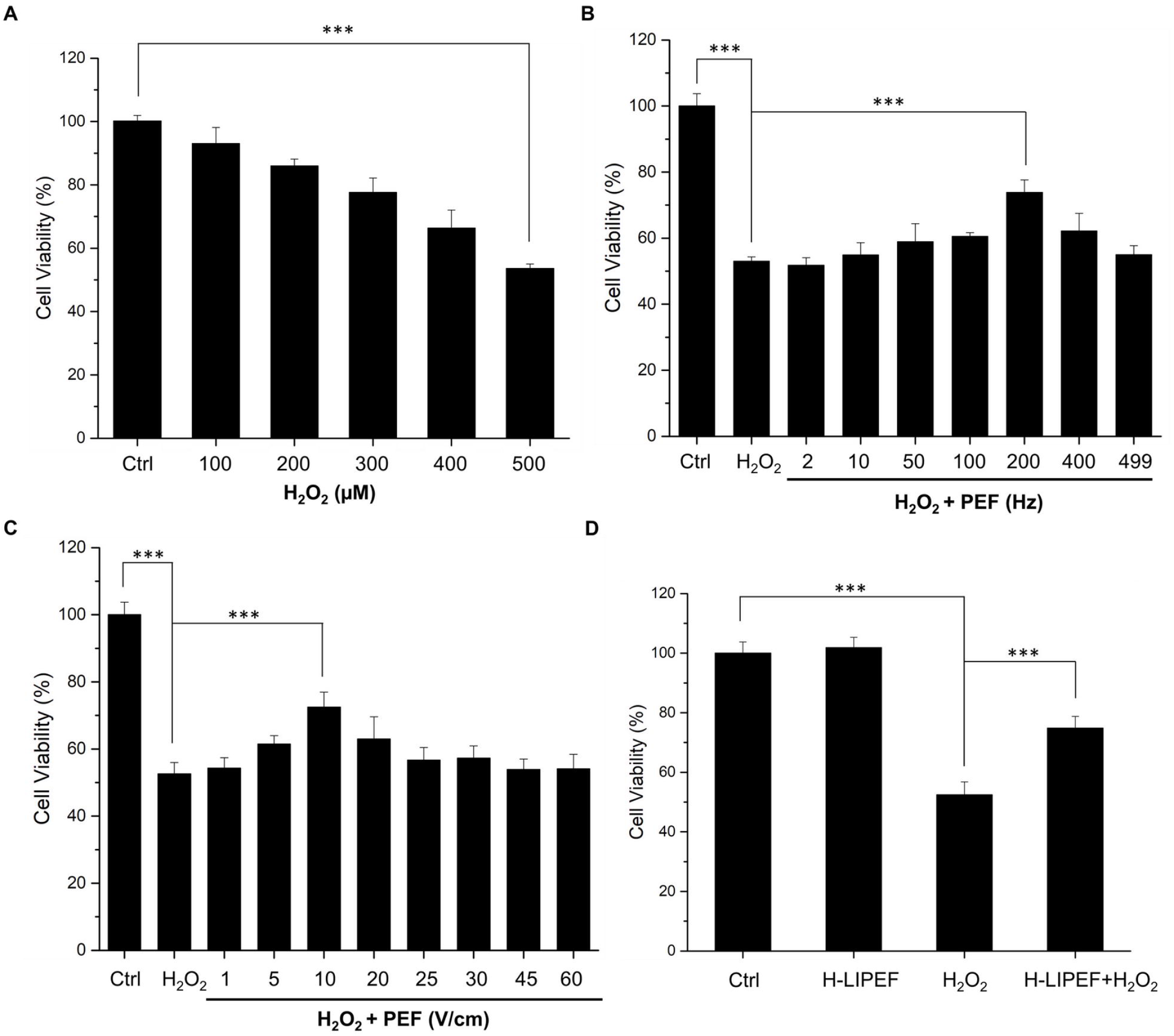
Effect of H-LIPEF on H_2_O_2_-induced cytotoxicity in SH-SY5Y cells. (A) The concentration response curve of SH-SY5Y cells treated with different concentrations of H_2_O_2_ and the cell viability was measured by MTT assay at 24 h after the H_2_O_2_ treatment. (B) SH-SY5Y cells were pretreated with 10 V/cm PEF at different pulse frequencies and challenged with 500 µM H_2_O_2_ for 24 h and then the cell viability was measured by MTT assay. (c) SH-SY5Y cells were pretreated using 200 Hz PEF with different field intensities and challenged with 500 µM H_2_O_2_ for 24 h and then the cell viability was measured by MTT assay. (D) SH-SY5Y cells were pretreated using 200 Hz H-LIPEF with field intensity of 10 V/cm and challenged with or without 500 µM H_2_O_2._ The cell viability was also measured by MTT assay at 24 h after the H_2_O_2_ treatment. Data represent the mean ± standard deviation (n=3). The concentration response curve of SH-SY5Y cells was statistically analyzed using one-way ANOVA with Tukey’s post hoc test (^***^P < 0.001). Comparisons of the effect of H_2_O_2_ and H-LIPEF on the cell viability were analyzed using two-way ANOVA with Tukey’s post hoc test (^***^P < 0.001).

**Fig 3.**
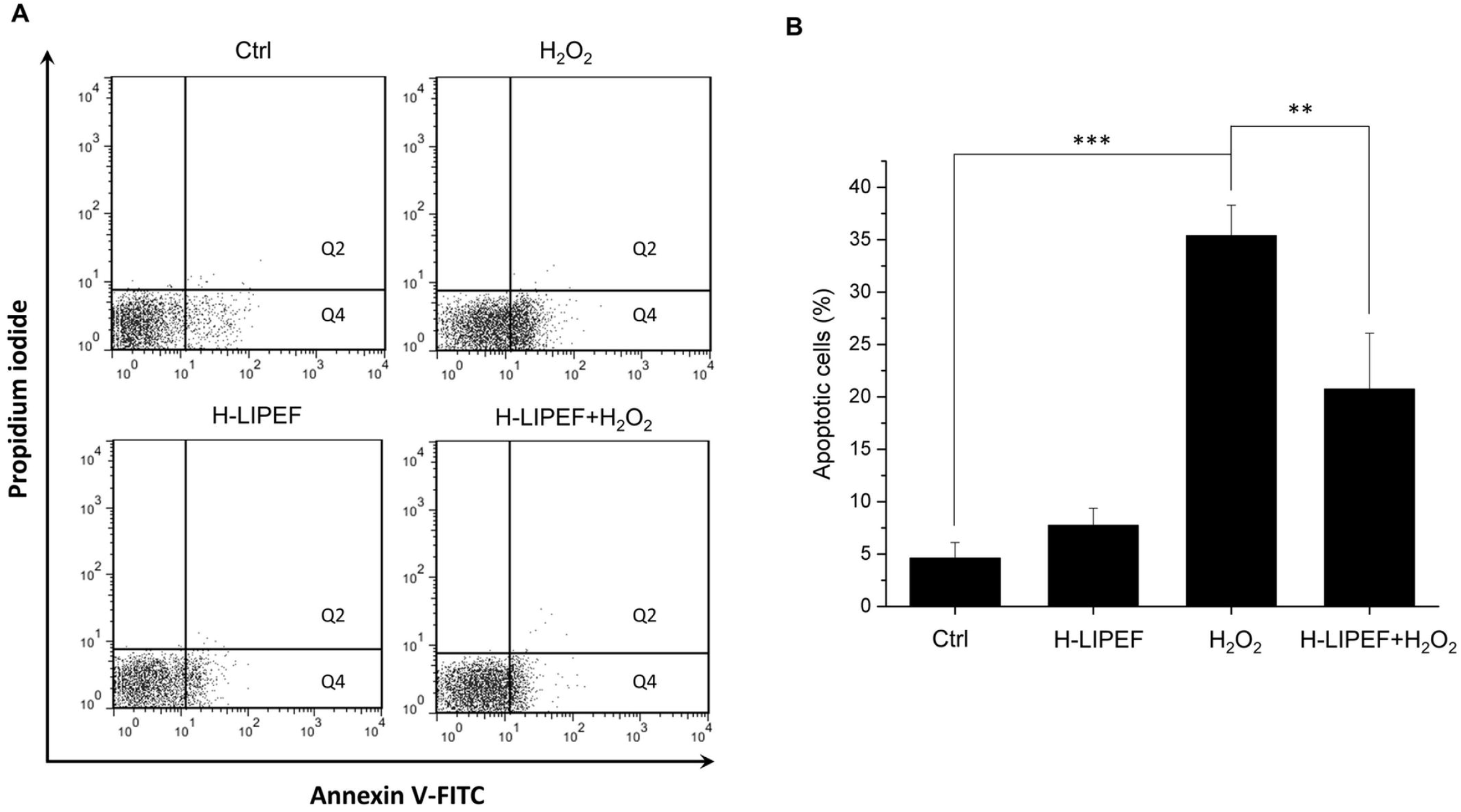
Assessment of the mode of cell death via Annexin V-FITC/PI double staining. (A) SH-SY5Y cells were pretreated using 200 Hz H-LIPEF with field intensity of 10 V/cm and challenged with or without 500 µM H_2_O_2_ for 24 h. The apoptotic cells were then assessed by flow cytometric analysis of Annexin V-FITC and PI staining. The cells in the Q2 and Q4 quadrants indicate Annexin V-positive apoptotic cells. (B) Quantification of SH-SY5Y apoptotic cells. Comparisons of the effect of H_2_O_2_ and H-LIPEF on the apoptotic cells were analyzed using two-way ANOVA with Tukey’s post hoc test (^**^P < 0.001 and ^**^P < 0.01).

**Fig 4.**
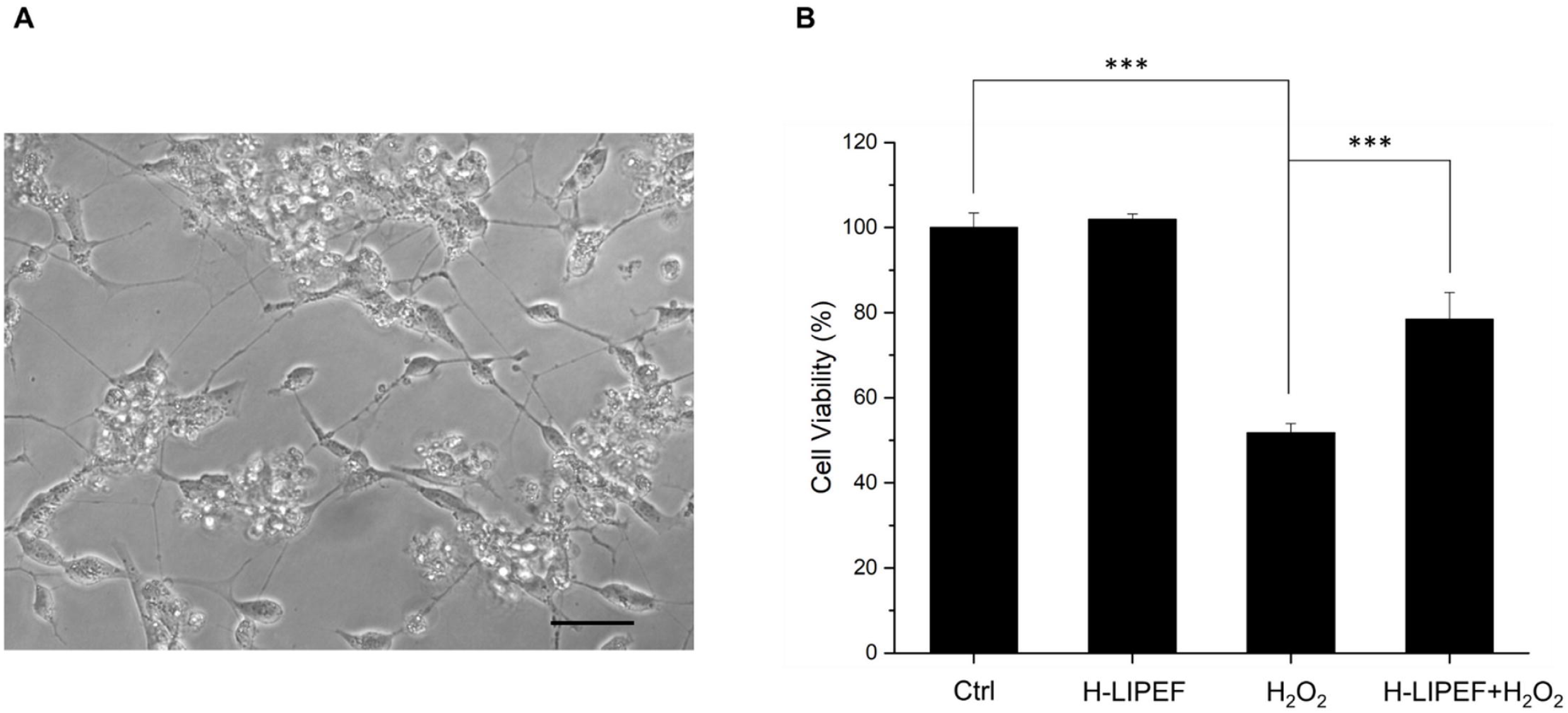
Effect of H-LIPEF on H_2_O_2_-induced cytotoxicity in differentiated SH-SY5Y cells. (A) Morphology of differentiated SH-SY5Y cells. Scale bar = 50 µm. (B) Differentiated SH-SY5Y cells were pretreated using 200 Hz H-LIPEF with field intensity of 10 V/cm and challenged with or without 500 µM H_2_O_2_ for 24 h. The cell viability was then determined by MTT assay. Comparisons of the effect of H_2_O_2_ and H-LIPEF on the cell viability were analyzed using two-way ANOVA with Tukey’s post hoc test (^***^P < 0.001).

### Effect of H-LIPEF exposure on Aβ-induced cytotoxicity in SH-SY5Y cells

For AD disease model, SH-SY5Y cells were treated with aggregated Aβ peptide, and the results showed that 25 or 50 µM Aβ treatment significantly reduced the viability of SH-SY5Y cells (Fig 5A). To confirm the Aβ aggregation state, the ThT dye was used to measure the β-sheet formation. The result showed that the fluorescence of the prepared Aβ stock solution was greatly higher than the ThT dye alone control group (Fig 5B), indicating that the Aβ stock solution was indeed in the β-sheet rich aggregated state. As shown in Fig 5C and 5D, the H-LIPEF treatment greatly improved the viability and restored the cell morphology of Aβ-treated SH-SY5Y cells, indicating the curative effect of H-LIPEF in AD disease model in vitro. The cell numbers were quantified using the ImageJ software [18] to investigate the effect of H-LIPEF on Aβ-induced cell number loss. The results showed that H-LIPEF retained the cell numbers of Aβ-treated SH-SY5Y cells (Fig 5E).

**Fig 5.**
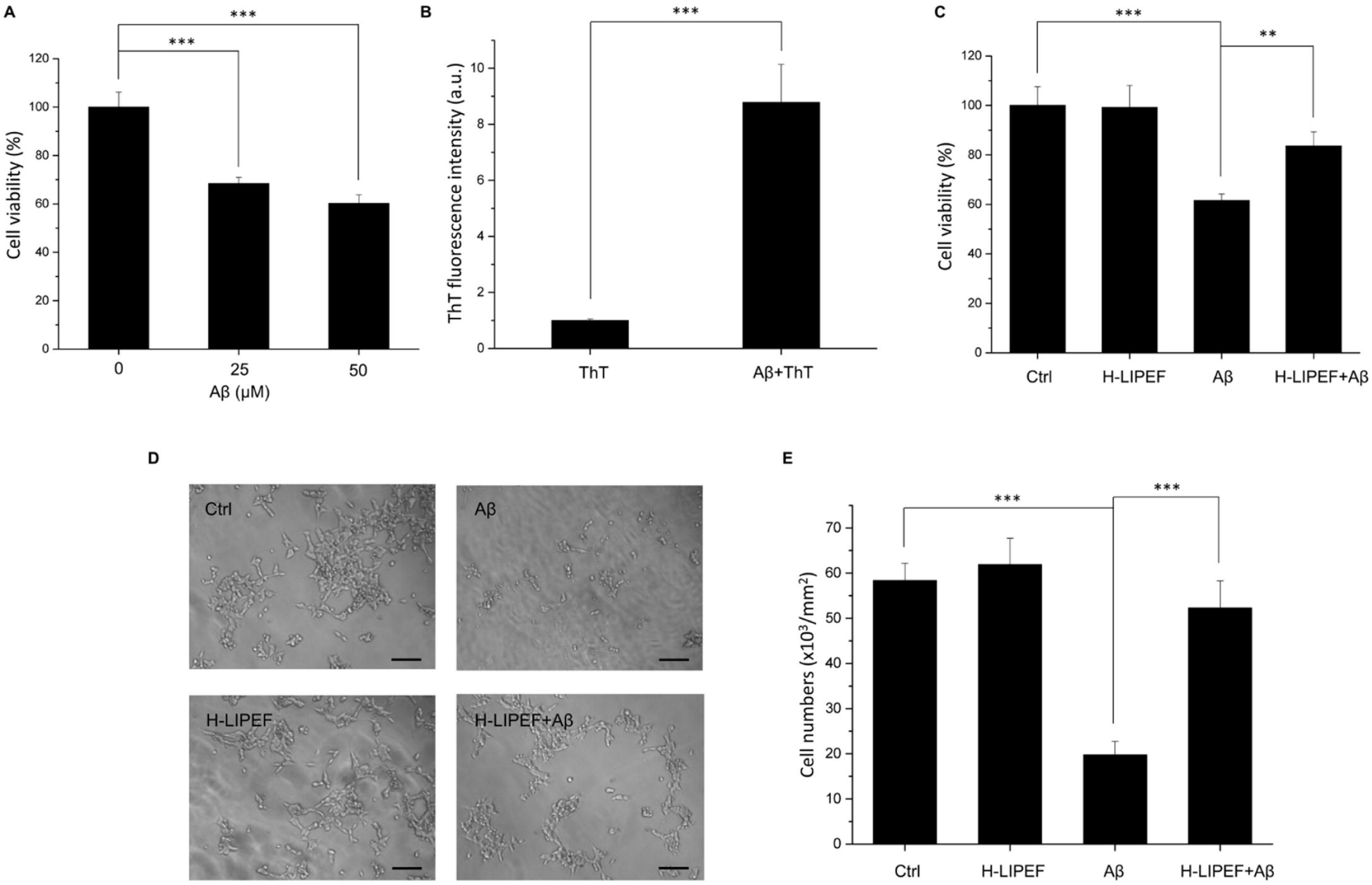
Effect of H-LIPEF on Aβ-induced cytotoxicity in SH-SY5Y cells. (A) The cell viability of SH-SY5Y cells treated with 25 or 50 µM Aβ for 4 days. (B) The Aβ aggregation state was assessed by the ThT fluorescence. (C) The SH-SY5Y cells were pretreated with H-LIPEF for 4 h and then exposed to 50 µM Aβ in the continuous administration of H-LIPEF for another 4 days, and the cell viability was measured by MTT assay. (D) Representative light microscopy images of SH-SY5Y cells after Aβ and/or H-LIPEF treatment. The integrity of the cells was destructed by Aβ, and the H-LIPEF treatment caused the protective effect and retained the cell morphology. Scale bar = 100 µm. (E) Quantification of the cell numbers in the light microscopy images by ImageJ software. Data represent the mean ± standard deviation (n=3). Cell viability of SH-SY5Y cells treated with different concentration of Aβ and the ThT fluorescence experiment were statistically analyzed using one-way ANOVA with Tukey’s post hoc test (^***^P < 0.001). Comparisons of the effect of Aβ and H-LIPEF on the cell viability and the cell numbers were analyzed using two-way ANOVA with Tukey’s post hoc test (^***^P < 0.001 and ^**^P < 0.01).

### H-LIPEF attenuates H_2_O_2_-induced ROS generation

Since the onset of AD is associated with ROS accumulation, we further examined the effect of H-LIPEF on the ROS level regulation by flow cytometry (Fig 6A). Quantification results (Fig 6B) showed that the ROS level was increased to 175% of the control level after exposure to 500 µM H_2_O_2_ for 24 h. Compared to the group of H_2_O_2_ alone, the H-LIPEF treatment greatly ameliorated the H_2_O_2_-induced ROS level elevation, indicating that H-LIPEF could have the ability to decrease the ROS level.

**Fig 6.**
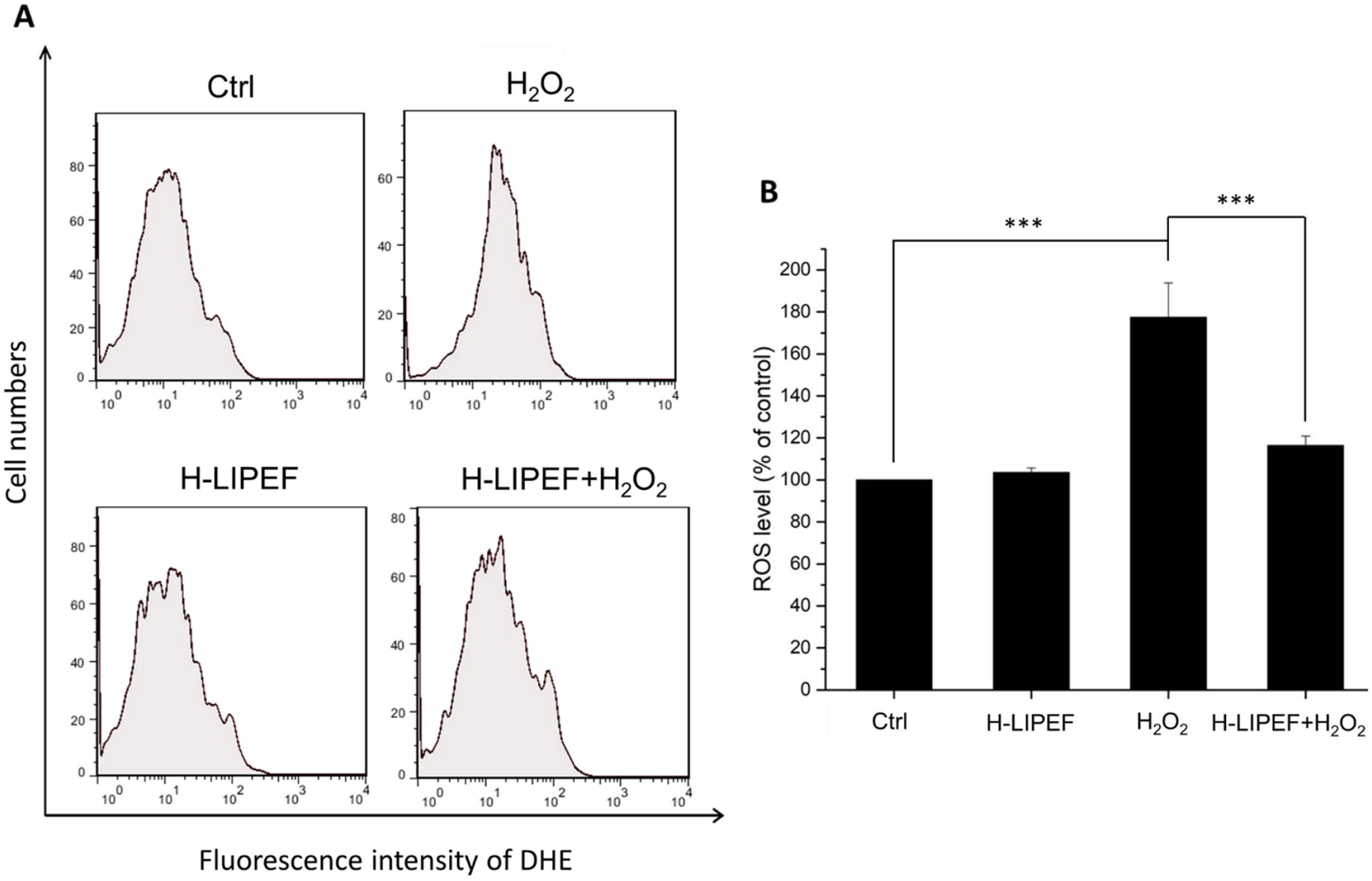
Effect of H-LIPEF on H_2_O_2_-induced ROS generation in SH-SY5Y cells. (A) ROS level was measured by flow cytometry with DHE fluorescent dye. (B) Quantification of the ROS levels after H-LIPEF, H_2_O_2_, or H-LIPEF+H_2_O_2_ treatment. Data represent the mean ± standard deviation (n=3). Comparisons of the effect of H_2_O_2_ and H-LIPEF on the ROS generation were analyzed using two-way ANOVA with Tukey’s post hoc test (^***^P < 0.001).

### H-LIPEF attenuates MMP loss in H_2_O_2_-treated cells

Substantial evidence suggests that mitochondria play important roles in NDDs through the accumulation of net cytosolic ROS production [19]. In healthy cells, the levels of MMP are kept relatively stable, and the depletion of MMP suggests the loss of mitochondrial membrane integrity. Apoptotic signals are usually initiated by such mitochondrial dysfunctions. To examine whether the neuroprotective effect of H-LIPEF is associated with mitochondrial functions, the MMP is further analyzed by the flow cytometry with DiOC_6_(3) fluorescent dye. As shown in Fig 7A, it was found that H_2_O_2_ caused mitochondrial dysfunction, decreasing the MMP and therefore decreased the intensity of the DiOC_6_(3) fluorescent signal while H-LIPEF alone did not change the MMP. From the quantification results (Fig 7B), we found that the cells with decreased MMP drastically increased after the H_2_O_2_ treatment, and H-LIPEF pretreatment noticeably suppressed the H_2_O_2_-induced dissipation of MMP, so the cells with decreased MMP reduced significantly. The results indicate that H-LIPEF could prevent the mitochondria from damaging by the oxidative stress and thus caused the blockage of the apoptotic signaling transduction.

**Fig 7.**
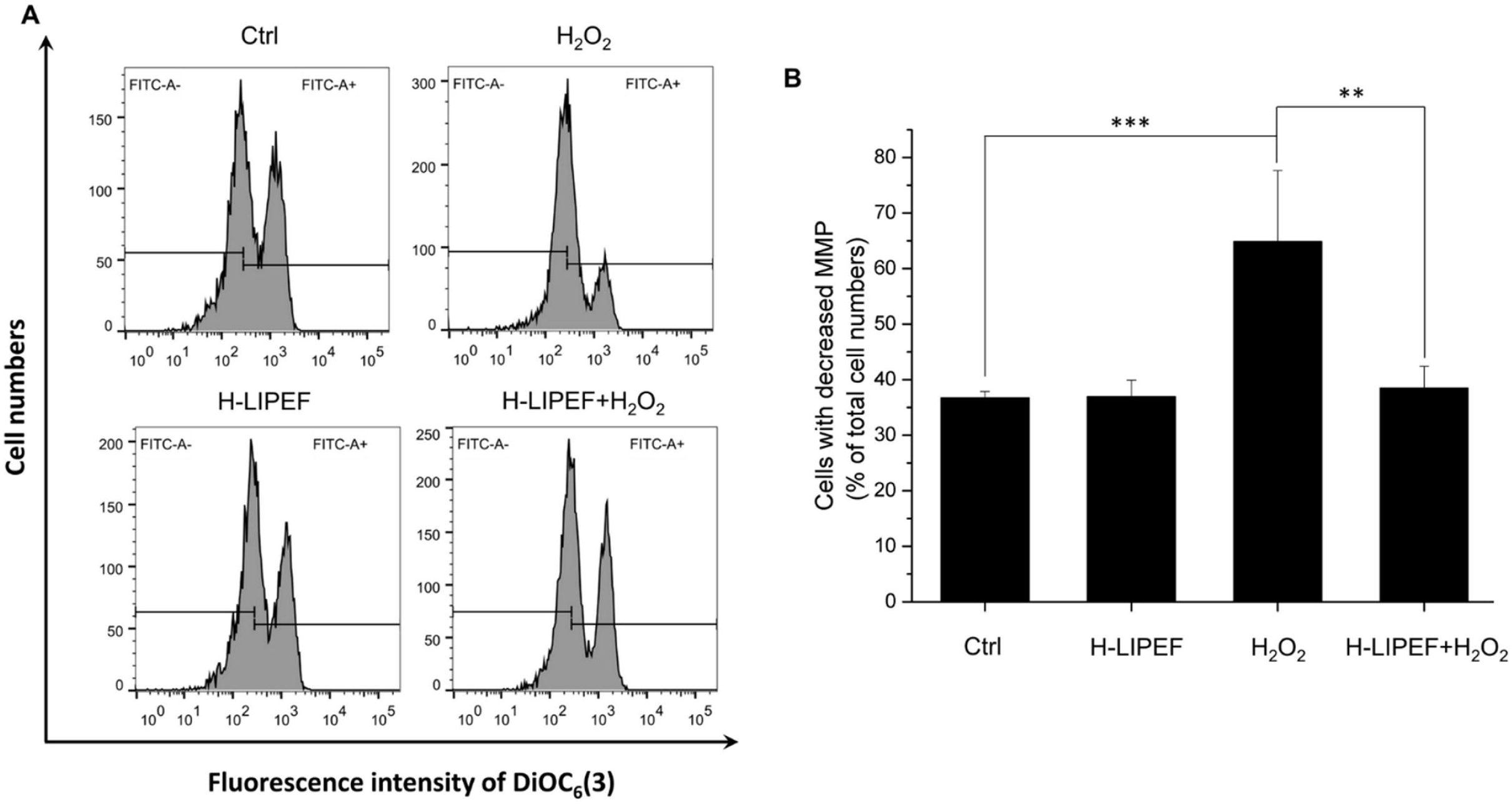
Effect of H-LIPEF on H_2_O_2_-induced MMP reduction in SH-SY5Y cells. (A) MMP was analyzed by flow cytometry with DiOC_6_(3) fluorescent dye. (B) Quantification of the cells with decreased MMP after H-LIPEF, H_2_O_2_, or H-LIPEF+H_2_O_2_ treatment. Data represent the mean ± standard deviation (n=3). Comparisons of the effect of H_2_O_2_ and H-LIPEF on the MMP were analyzed using two-way ANOVA with Tukey’s post hoc test (^***^P < 0.001 and ^**^P < 0.01).

### Effect of H-LIPEF treatment on ROCK protein expression in SH-SY5Y cells

In our previous study, the neuroprotective effect of LIPEF was found to be associated with the inhibition of the H_2_O_2_-induced activation of ROCK pathway in the motor neuron cell line NSC-34 [12]. In this work, we examined the ROCK protein expressions in order to investigate whether the protective effect of high frequency LIPEF on human neuroblastoma SH-SY5Y cells was related to the ROCK pathway. As shown in Fig 8, the expression of ROCK protein is slightly higher after H_2_O_2_ treatment but almost unaffected after the treatment of both H-LIPEF and H_2_O_2_. This phenomenon points out that although the H-LIPEF treatment can protect SH-SY5Y cells, the main mechanism is not associated with the ROCK pathway. Therefore, other prosurvival protein pathways should be considered in the following experiments.

**Fig 8.**
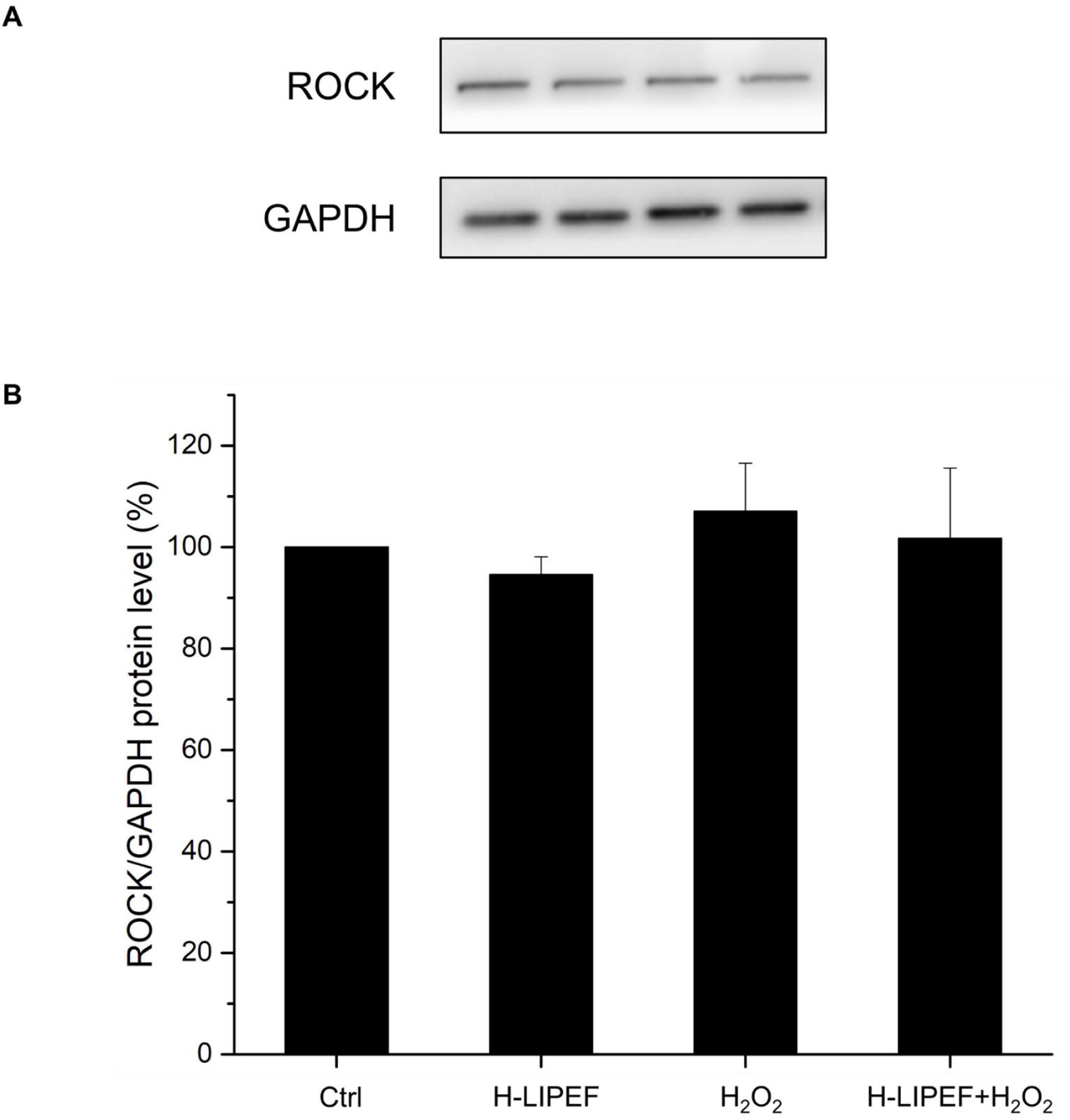
Effect of H-LIPEF on expressions of ROCK protein in SH-SY5Y cells. (A) Western blot analysis of ROCK protein expressions. (B) Quantification of ROCK expression level after H-LIPEF, H_2_O_2_, or H-LIPEF+H_2_O_2_ treatment. The expression levels were normalized to GAPDH and each relative expression level was compared with control. Data represent the mean ± standard deviation (n=3).

### Effect of H-LIPEF exposure on ERK-Nrf2/CREB signalling pathways in H_2_O_2_-treated SH-SY5Y cells

We then studied the signalling pathways that could participate in the neuroprotective mechanism of H-LIPEF stimulation. ERK is one of the major cascades of mitogen-activated protein kinases (MAPK) family and has been known to be widely associated with cell survival [20]. Upon phosphorylation, the active ERK proteins enter into the nucleus where they can phosphorylate certain transcription factors, thus regulating the expression of specific genes that contribute to cell survival [21]. Therefore, in our study, we examined the phosphorylation of ERK and the downstream transcription factors including cAMP response element binding protein (CREB) and nuclear factor erythroid 2-related factor 2 (Nrf2) by western blot analysis.

In comparison to the untreated control cells, as shown in Fig 9A, the normalized level of p-ERK (p-ERK/t-ERK) was found to slightly increase in the cells exposed to H-LIPEF alone for 24 h. In contrast, the normalized level of p-ERK (p-ERK/t-ERK) in the cells was significantly decreased when exposed to 500 µM H_2_O_2_ for 24 h without the treatment of H-LIPEF. In addition, the result showed that the H-LIPEF treatment induced a significant recovery of the p-ERK and t-ERK levels in the cells exposed to 500 µM H_2_O_2_. The results indicate that the neuroprotection provided by H-LIPEF could be related to the recovery of the H_2_O_2_-induced decline of ERK expression and its phosphorylation. In the following, we further examined the expression levels of Nrf2 and p-CREB proteins using western blot analysis. Nrf2 is a key transcription factor that regulates the expression of antioxidant proteins participating in the protection of oxidative damage triggered by injury and inflammation. As shown in Fig 9B, the level of Nrf2 was also increased in H-LIPEF treated SH-SY5Y cells compared to the untreated cells. In contrast, the level of Nrf2 in the cells exposed to 500 µM H_2_O_2_ for 24 h but without the treatment of H-LIPEF was decreased. Meanwhile, the H-LIPEF treatment was found to recover the expression of Nrf2 in the cells exposed to 500 µM H_2_O_2_ obviously. This result showed that the neuroprotective effect was due in part to the H-LIPEF induced Nrf2 expression and the antioxidant effect. Next, CREB is also an important transcription factor triggering the expression of prosurvival proteins. As shown in Fig 9C, the levels of both p-CREB and t-CREB in cells were greatly decreased when exposed to 500 µM H_2_O_2_ for 24 h without the treatment of H-LIPEF. We further found that the normalized level of p-CREB (p-CREB/t-CREB) was significantly reduced upon H_2_O_2_ treatment while H-LIPEF restored the p-CREB/t-CREB level. The results indicate that the neuroprotection provided by H-LIPEF could be related to the recovery of the H_2_O_2_-induced decline of CREB expression and its phosphorylation. Furthermore, it is widely known that CREB phosphorylation regulates Bcl-2 family proteins including anti-apoptotic Bcl-2 and pro-apoptotic Bax proteins, and the ratio of Bcl-2/Bax determines whether neurons undergo survival or death after an apoptotic stimulus. Therefore, the expressions of Bcl-2 and Bax protein levels were also examined in the study. As shown in Fig 9D, for SH-SY5Y cells under the H-LIPEF treatment alone, the expression levels of Bcl-2 and Bax were increased and decreased respectively, leading to a higher elevation of the Bcl-2/Bax ratio compared to the untreated control cells. Meanwhile, in comparison to the untreated control cells, the level of Bcl-2/Bax ratio of SH-SY5Y cells was found to decrease when exposed to 500 µM H_2_O_2_, indicating that the cells had undergone apoptosis. Moreover, it was observed that the H-LIPEF treatment notably restored the expression of Bcl-2/Bax ratio in the cells exposed to 500 µM H_2_O_2_. To sum up, the results showed that the H-LIPEF treatment induced the anti-apoptotic effect by regulating the Bcl-2 family proteins via modulating the ERK/CREB pathways and thus promoted the cell survival against oxidative stress.

**Fig 9.**
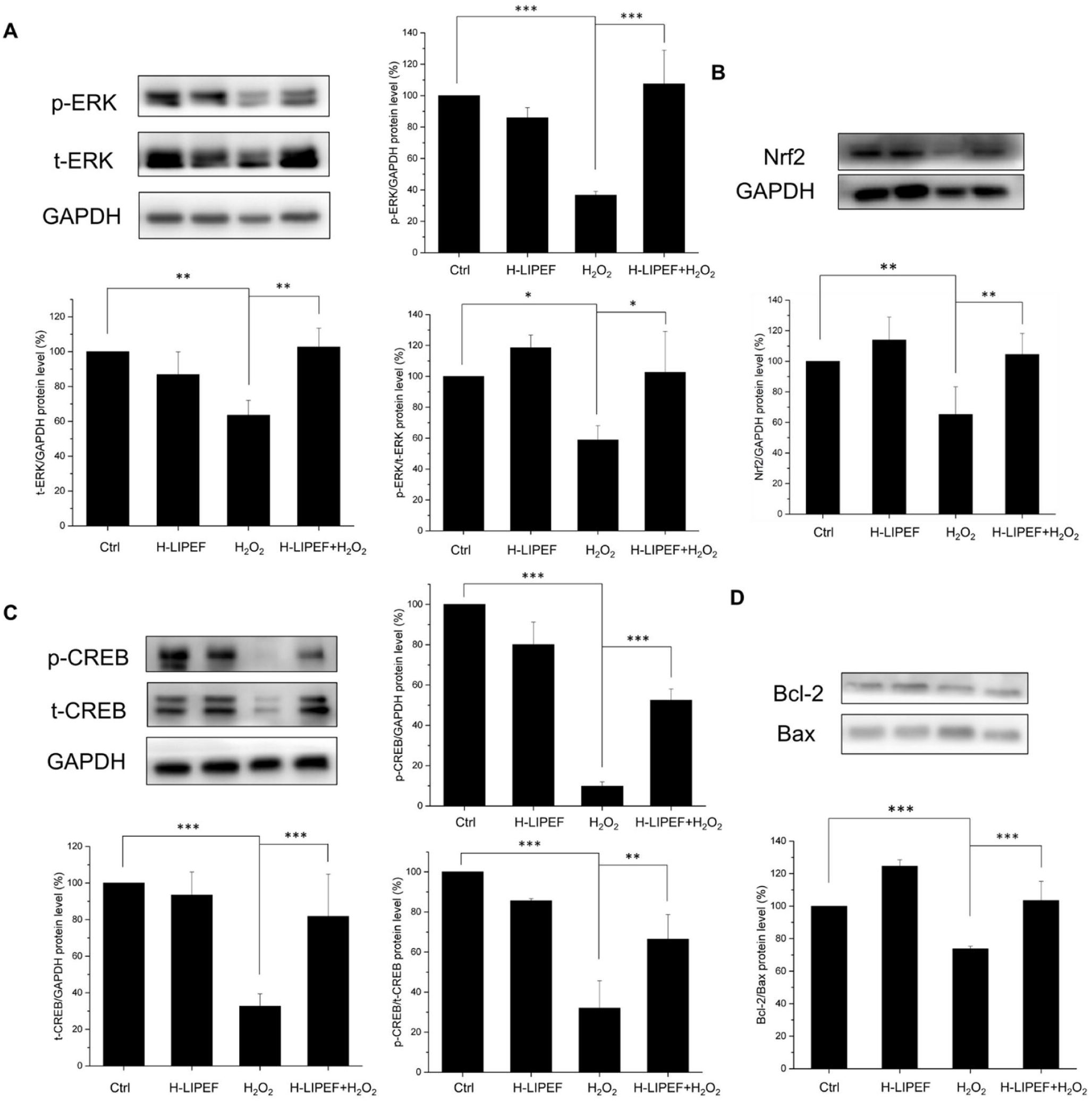
Effect of H-LIPEF on ERK/Nrf2 and ERK/CREB signalling pathways and related protein expressions. Western blot analysis of p-ERK and t-ERK (A), Nrf2 (B), p-CREB and t-CREB (C), Bcl-2 and Bax (D) protein expressions. The expression levels of p-ERK and p-CREB were normalized to t-ERK and t-CREB, respectively. The expression level of Nrf2 was normalized to GAPDH and each relative expression level was compared with control. Data represent the mean ± standard deviation (n=3). Comparisons of the effect of H_2_O_2_ and H-LIPEF on different protein levels were analyzed using two-way ANOVA with Tukey’s post hoc test (^***^P < 0.001, ^**^P < 0.01 and ^*^P < 0.05).

### Effect of PD98059 on the H-LIPEF activated ERK pathway and protective effect

To confirm the neuroprotective effect of H-LIPEF is related to the ERK pathway, we inhibited the MAPK kinases MEK1 and MEK2, the upstream kinases of ERK, by the inhibitor PD98059 [22] in the study. In Fig 10A, we found that H-LIPEF could significantly increase the cell viability of SH-SY5Y cells to about 80% of the control level under H_2_O_2_ stress. In the presence of MEK inhibitor, the p-ERK level was indeed reversed and the neuroprotective effect of H-LIPEF treatment against H_2_O_2_-induced cell apoptosis was abrogated (Fig 10A and 10B). The inhibitor PD98059 greatly inhibited the phosphorylation of ERK protein, and therefore resulted in the accumulation of total ERK which was in agreement with previous studies [23, 24]. The results indicate that the ERK pathway was involved to mediate the neuroprotective effect of H-LIPEF treatment against H_2_O_2_ toxicity. On the whole, the data point out that H-LIPEF treatment may produce neuroprotective effect via activation of the ERK signalling pathway in the H_2_O_2_-treated SH-SY5Y human neural cells.

**Fig 10.**
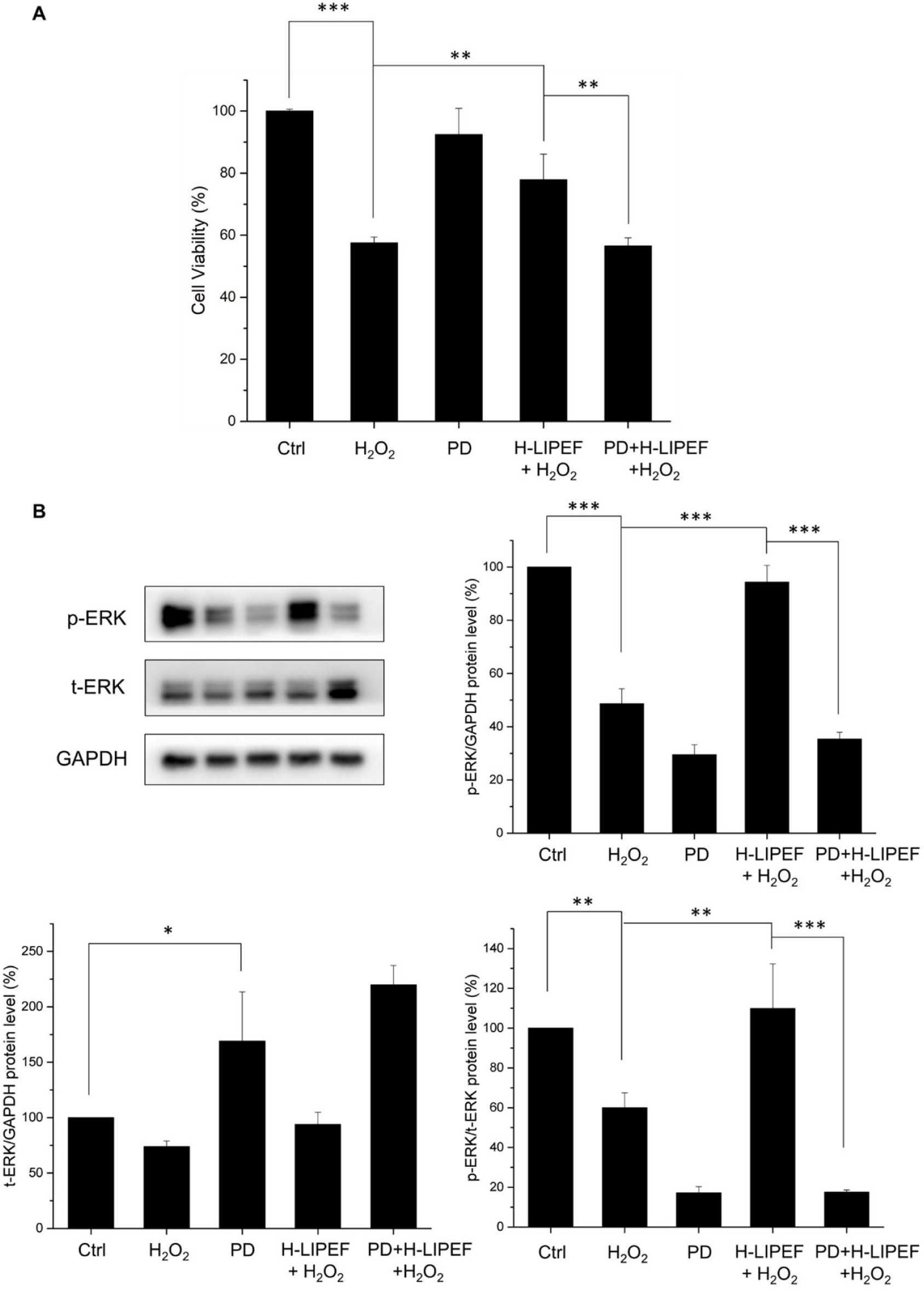
Effect of PD98059 on the neuroprotective effect of H-LIPEF and p-ERK protein expression induced by H-LIPEF stimulation. (A) H-LIPEF increased the cell viability of SH-SY5Y cells under H_2_O_2_ oxidative stress. The neuroprotective effect of H-LIPEF treatment was abrogated by addition of the MEK inhibitor PD98059. (B) The western blot analysis showed that the inhibitor PD98059 reversed the activated level of p-ERK induced by H-LIPEF treatment. The p-ERK expression levels were normalized to t-ERK. We used the abbreviation “PD” to represent the MEK inhibitor PD98059 in the figure. Data represent the mean ± standard deviation (n=3). Comparisons of the effect of PD on the neuroprotective effect of H-LIPEF and p-ERK protein levels were analyzed using two-way ANOVA with Tukey’s post hoc test (^***^P < 0.001 and ^**^P < 0.01).

## Discussion

The focus of this study was to probe the neuroprotective effect of H-LIPEF on SH-SY5Y cells. The results demonstrated that application of high-frequency (∼200 Hz) and low-intensity (∼10 V/cm) pulsed electric field produced the most significant protection for SH-SY5Y human neural cells against H_2_O_2_- and Aβ-induced apoptosis. The western blot results exhibited that H-LIPEF triggered the ERK signalling pathway and mediated the activation of CREB and Nrf2 transcription factors.

As the most common form of age-related NDDs, AD features excessive aggregation of Aβ and intracellular microtubule-associated protein tau, known as neurofibrillary tangles. With many questions remaining unanswered, some researchers believe that the amyloid cascade hypothesis offers the best explanation for the cause of AD [25], maintaining that Aβ is the causative factor of Alzheimer’s pathology and that the neurofibrillary tangles, cell loss, vascular damage, and dementia all result directly from such deposition. Although the mechanism of Aβ-induced cytotoxicity has yet to be elucidated completely, several studies have proven that the oxidative stress plays a vital role in Aβ-induced apoptosis [26-28]. Therefore, researchers have been striving to find various antioxidants that can protect against oxidative stress and alleviate the symptoms of NDDs [29-31]. However, the therapeutic effect of most antioxidant compounds is limited, due to their incapability to cross the BBB. This is reflected in the failure of antioxidant therapies in clinical trial [6, 7, 10]. Even for the fat soluble Vitamin E, there have been no promising results as far as we know [8, 9]. Therefore, an effective AD treatment strategy is urgently needed. Among different ROS species, H_2_O_2_ is one of the molecules most related to the progression of AD. In the study, H_2_O_2_ was applied as the oxidative stress which caused cell death at certain dosage. The results found that H_2_O_2_ treatment increased the ROS level and H-LIPEF treatment significantly ameliorated the H_2_O_2_-induced ROS level elevation, underscoring the antioxidant ability of H-LIPEF. The study employed MTT and MMP assays to assess the cell viability and mitochondria integrity. The findings in both results showed that H-LIPEF provided the neuroprotective effect and increased the viability and MMP in the H_2_O_2_-treated neural cells. Furthermore, Aβ was used to investigate the neuroprotective effect of H-LIPEF in the AD disease model. The MTT and light microscopy results also showed that H-LIPEF greatly improved the viability and retained the cell integrity in the Aβ-treated cells. As demonstrated by the recent studies in cancer treatment [32, 33], electric fields can penetrate the human brain without being restricted by BBB, which suggests that H-LIPEF may be an alternative therapeutic mode in NDDS treatment.

The electric stimulation has a long history of medical applications, including wound healing [34], muscle relaxation [35], and increase of blood circulation [36], often involving passage of electric current through the injury parts via direct contact between the body and implanted electrodes. However, the invasive or contact manner significantly dampens the feasibility of the treatment or even causes harms to cells or body, such as electrolysis, burns and muscle spasms [37]. Therefore, it is desirable for electrotherapy to be carried out in a non-invasive or non-contact manner. Our previous study demonstrated a non-contact electric stimulation device [13], which operates as a parallel-plate capacitor generating electric field between two parallel electrodes (Fig 1A). Therefore, the electric field with such capacitive coupling mode can pass through biological samples to achieve specific biological effect without current actually flowing through cells or media via contact with electrodes. It is noteworthy that the strength of electric field passing through culture dishes or media is the

same as the voltage for electrodes. Besides, the electric field waveform employed was a PEF, which is characterized by its multi-frequency component nature. Pulse train signal is composed of multiple sinusoidal subcomponents with different frequencies and intensities [12]. When PEF enters a biological sample, it encounters different proteins, cells and tissues with different dielectric constants and is therefore dispersed to multiple frequencies electrically. Multi-frequency sinusoidal waves may interact with molecules, proteins, or organelles simultaneously to induce specific bioeffect in the PEF exposure. In our previous studies, the non-contact LIPEF exhibited promising potential in activating anticancer agent in pancreatic cancer treatment [38] and also in the neuroprotection of mouse motor neurons [12]. In the current study, LIPEF with different parameters was applied to the human neural cells for examination of its neuroprotective effect. Unexpectedly, the LIPEF parameter applied in mouse motor neurons NSC-34 turned out to be ineffective for human SH-SY5Y cells. The study found that for human SH-SY5Y cells, the neuroprotective effect of LIPEF was more pronounced in the high-frequency region, especially near 200 Hz, quite different from the parameter at 2 Hz for NSC-34 mouse cell line. For higher-frequency LIPEF, it consists more high-frequency components in its composition. Therefore, the neuroprotective effect of H-LIPEF for human SH-SY5Y neural cells may be attributable to the employment of high-frequency components participating in the treatment. The results also showed that the therapeutic window of the H-LIPEF intensity for SH-SY5Y cells was near 10 V/cm, which was smaller than the most effective parameter (60 V/cm) for NSC-34 cells. Therefore, it is evident that the pulse frequency and the field intensity are critical parameters for therapeutic applications. The parameters (frequency, intensity, and pulse duration) of LIPEF need to be fine-tuned and optimized for different types of cells or diseases. Furthermore, the study proposes combination of multiple pulses with different parameters to treat the targeted region simultaneously.

As for the neuroprotective mechanism, it was found previously that the LIPEF treatment obviously suppressed the H_2_O_2_-enhanced expression of ROCK protein in NSC-34 cells. However, for SH-SY5Y cells, the H-LIPEF simulation did not have significant impact on ROCK protein expression. It is thus known that the neuroprotective mechanism is not through the ROCK pathway. In the western blot results, the study proved that such neuroprotective effect induced by H-LIPEF was through activation of ERK pathway. The ERK pathway plays an important role in regulating of proliferation and differentiation as well as survival of various cell types [39]. It transmits external cues, including growth factors into signalling cascade via the phosphorylation of ERK. Several studies have suggested that the stimulation of ERK pathway may be a potential way for neuroprotection [20, 21]. Here, the study showed that the H-LIPEF stimulation induced a significant recovery of the p-ERK level in the cells under H_2_O_2_ oxidative stress. The result indicates that the neuroprotection effect produced by H-LIPEF could be related to the recovery of the H_2_O_2_-induced reduction of p-ERK. Moreover, it should be noted that the normalized p-ERK level (p-ERK/t-ERK) stimulated by H-LIPEF alone was only slightly higher than the untreated control cells without significant difference (Fig 9A). The study found that H-LIPEF did not cause cell proliferation as assessed by MTT assay (Fig 2D). Therefore, the result confirmed that the neuroprotective effect of H-LIPEF stimulation was not caused by the increased cell numbers, but by specific protein pathways induced by H-LIPEF on SH-SY5Y cells. Furthermore, since the ERK pathway is initiated by the binding of epidermal growth factor (EGF) to the epidermal growth factor receptor (EGFR), we speculate that H-LIPEF could have an interaction with EGFR and cause the conformation change of EGFR, thus triggering the ERK pathway. Previous study had shown that magnetic field could alter the conformation of TRPV1 ion channel, making it more favorable for capsaicin binding [40]. The results implicate that the PEF or magnetic field with different parameters could activate different signalling pathways. In the study, H-LIPEF with 200 Hz frequency and 10 V/cm field intensity was found to significantly recover the p-ERK protein level and produce the neuroprotective effect. The phosphorylation of ERK protein could result in the activation of its kinase activity and lead to phosphorylation of many downstream targets including certain transcription factors containing CREB and Nrf2 required for neuroprotection.

CREB is an important transcription factor activated by phosphorylation from ERK or other kinases. It triggers expression of neurotrophins and antiapoptotic proteins such as brain-derived neurotrophic factor (BDNF) and B-cell lymphoma 2 (Bcl-2) [41]. Besides, CREB downregulation is related to the pathology of AD [42], and thus increasing the CREB expression has been considered as a potential therapeutic strategy for AD. In this study, the result found that the H-LIPEF treatment notably recovered the expression of p-CREB in the cells exposed to 500 µM H_2_O_2_. On one hand, it is known that CREB phosphorylation regulates Bcl-2 family proteins including anti-apoptotic Bcl-2 and pro-apoptotic Bax proteins. The antiapoptotic protein, Bcl-2, has been proposed to be a pivotal component for neuroprotection [43]. It suppresses apoptosis by heterodimerizing with Bax protein and neutralizes the effects of Bax. When Bcl-2 is present in excess, cells are protected against apoptosis. It should be also pointed out that when Bax is excessive and the homodimers of Bax dominate, cells are susceptible to programed cell death. Therefore, it is the ratio of Bcl-2 to Bax which determines the fate of a cell [44]. In the results, we found that the H-LIPEF treatment obviously restored the expression of Bcl-2/Bax ratio in the cells exposed to 500 µM H_2_O_2_, indicating the prosurvival effect of H-LIPEF in SH-SY5Y cells.

Another possible mechanism for neuroprotection of the H-LIPEF stimulation may be associated with the Nrf2 protein. Nrf2 is also an important transcription factor activated by ERK pathway. It regulates the expression of various antioxidant proteins and maintains their redox states in mammalian cells [45]. The results showed that the H-LIPEF treatment obviously recovered the expression of Nrf2 in the cells exposed to 500 µM H_2_O_2_, indicating that the neuroprotective effect was due in part to the H-LIPEF induced Nrf2 expression and the antioxidant effect.

In the study, our results confirmed the neuroprotective effect of H-LIPEF stimulation on SH-SY5Y cells. Because electric field can be modified to focus on a targeted region, it affirms the applicability of H-LIPEF treatment for different human organs. Moreover, it found that H-LIPEF technique can significantly reduce treatment risk and time, thanks to its non-contact and multi-frequency nature. In addition, the study also demonstrated the ability of H-LIPEF in rescuing the Aβ-induced cytotoxicity. Since the accumulation of misfolded proteins in brain is also a typical symptom of some NDDs, the study could shed light on future exploration for optimal LIPEF conditions in the treatment of other NDDs.

In summary, the study demonstrates for the first time that the physical stimulation by H-LIPEF can protect SH-SY5Y human neural cells from H_2_O_2_- and Aβ-induced cytotoxicity. The underlying molecular mechanism for the neuroprotective effect is partially attributed to the activation of ERK pathway and the function of some downstream proteins, such as CREB, Nrf2, and Bcl-2. Further studies are necessary to ascertain the optimal LIPEF conditions for other NDDs and the involvement of other proteins in the neuroprotective effect. The combinations of H-LIPEF with NDD drugs or natural compounds are also advisable for future studies.

## Acknowledgments

The authors would like to acknowledge the service provided by the Research Core Facilities 3 Laboratory of the Department of Medical Research at National Taiwan University hospital for use of flow cytometry system.

## Author Contributions

**Conceptualization:** Chih-Yu Chao.

**Data Curation:** Wei-Ting Chen, Guan-Bo Lin, Yu-Yi Kuo, Chih-Yu Chao.

**Formal analysis:** Wei-Ting Chen, Guan-Bo Lin, Yu-Yi Kuo, Chih-Hsiung Hsieh, Chueh-Hsuan Lu, Yi-Kun Sun, Chih-Yu Chao.

**Funding acquisition:** Chih-Yu Chao.

**Investigation:** Wei-Ting Chen, Guan-Bo Lin, Yu-Yi Kuo, Chih-Yu Chao.

**Project Administration:** Chih-Yu Chao.

**Supervision:** Chih-Yu Chao.

**Validation:** Wei-Ting Chen, Guan-Bo Lin, Yu-Yi Kuo.

**Writing – original draft:** Wei-Ting Chen, Guan-Bo Lin, Chih-Yu Chao.

**Writing – review & editing:** Chih-Yu Chao.

